# The neurocomputational architecture of explore-exploit decision making

**DOI:** 10.1101/2021.06.21.449128

**Authors:** Jeremy Hogeveen, Teagan S. Mullins, John Romero, Elizabeth Eversole, Kimberly Rogge-Obando, Andrew R. Mayer, Vincent D. Costa

**Affiliations:** Department of Psychology & Psychology Clinical Neuroscience Center, Albuquerque, NM, USA; Department of Psychiatry & Behavioral Sciences, Albuquerque, NM, USA; Department of Neurology, University of New Mexico, Albuquerque, NM, USA; Mind Research Network/LBERI, Albuquerque, NM, USA; Department of Behavioral Neuroscience, Oregon Health and Science University, Portland, OR, USA

**Keywords:** Reward, explore-exploit, novelty-seeking, fMRI, computational modelling, frontopolar cortex

## Abstract

Humans and other animals often make the difficult decision to try new options (exploration) and forego immediate rewards (exploitation). Novelty-seeking is an adaptive solution to this explore-exploit dilemma, but our understanding of the neural computations supporting novelty-seeking in humans is limited. Here, we presented the same explore-exploit decision making task to monkeys and humans and found evidence that the computational basis for novelty-seeking is conserved across primate species. Critically, through computational model-based decomposition of event-related functional magnetic resonance imaging (fMRI) in humans, these findings reveal a previously unidentified cortico-subcortical architecture mediating explore-exploit behavior in humans.

## Introduction

The motivation to explore novel information instead of resorting to already learned behaviors is a bedrock for how humans learn across the lifespan. Yet, exploration comes at the cost of exploiting familiar options whose immediate consequences are known. Managing this tradeoff is referred to as the ‘explore-exploit’ dilemma, and these decisions are implemented via interactions between prefrontal and motivational brain regions (e.g. amygdala and striatum). However, considerable controversy remains regarding the neural computations that drive exploration. In one view, prefrontal cortex and motivational regions may work together to compute both the anticipated immediate and future value of choice options, driving the exploration of novel opportunities when it could lead to more rewards in the future (Costa and Averbeck, 2020a; Costa et al., 2019a; Wilson et al.). In another view, prefrontal cortex may override encoding of familiar outcomes in motivational regions while forming new decision policies (Choung et al., 2017; Daw et al., 2006; Domenech et al., 2020; Ebitz et al., 2018). Resolving the neurocomputational architecture of explore-exploit decisions could contribute to the development of more effective circuit-based treatments for transdiagnostic psychiatric challenges including inflexibility (Voon et al., 2017). Here, we probe the computations encoded in prefrontal and motivational regions during explore-exploit behavior in humans using model-based fMRI, and build a translational bridge for future invasive and noninvasive studies by demonstrating conservation of explore-exploit computations across humans and monkeys on the same task.

The neurocomputational bases of explore-exploit decisions in primates have been investigated using neural recordings in monkeys. A network comprising amygdala, ventral striatum, and orbitofrontal cortex encodes *both* the anticipated immediate *and* latent future value of choice options during explore-exploit decisions (Costa and Averbeck, 2020a; Costa et al., 2019a). However, these studies are restricted to *a priori* targeted recording sites and therefore limited in anatomical scope. Whole brain examinations of explore-exploit decisions via fMRI in humans have suggested that lateral frontopolar cortex (lFPC) and posterior parietal regions also play a role in explore-exploit decisions (Badre et al., 2012; Boorman et al., 2009; Daw et al., 2006). However, in these studies ‘exploration’ is typically defined as an overriding of ‘exploitative’ decision policies (i.e., failure to maximize value), making it difficult to determine whether FPC is encoding computations related to goal-directed exploration, random exploration as a function of decision noise, or poor reinforcement learning (Zajkowski et al., 2017). In the current study, human and monkey subjects performed a reinforcement learning task involving periodic insertion of novel stimuli with an unknown value. Novelty-seeking in this context is an evolved solution to the explore-exploit dilemma, which should explicitly signal a motivation to engage in goal-directed exploration. Importantly, recent advances in computational modelling with a normative agent framework enabled us to formally quantify the immediate and latent future value for each option, independent of observed behavior (Averbeck, 2015; Costa and Averbeck, 2020a; Costa et al., 2019a). To be clear, this allowed us to directly compare the value of exploring versus exploiting within the same brain regions—and across the whole brain.

## Results

We utilized a three-arm bandit task where novelty was used to motivate exploration (**Figure 1A**). Specifically, monkeys and humans performed speeded decisions between 3 neutral images assigned low (*p*_reward_=0.2), medium (*p*_reward_=0.5), or high (*p*_reward_=0.8) reward values. Periodically, a novel stimulus with a randomly assigned value was inserted, forcing participants to face a tradeoff between exploring the new option versus exploiting the best available alternative. Both monkeys and human participants were more likely to explore the novel stimulus on the early^1^ trials after a novel stimulus was introduced (Monkey: *M*=0.45, *SEM*=0.02; Human: *M*=0.46, *SEM*=0.02) than to exploit the best alternative option (Monkey: *M*=0.31, *SEM*=0.02, *t*_yuen_=3.27, *p*=0.02, *η*=0.88; Human: *M*=0.35, *SEM*=0.02, *t*_yuen_=2.96, *p*=0.007, *η*=0.55; **Figure 1B-C**). In turn, the tendency to exploit the best alternative option on early trials was higher than selection of the worst alternative (Monkey: *M*=0.24, *SEM*=0.01, *t*_yuen_=3.48, *p*=0.02, *η*=0.70; Human: *M*=0.19, *SEM*_human_=0.01, *t*_yuen_=3.93, *p*<0.001, *η*=0.79; **Figure 1B-C**). Looking at decision making over time, the probability of selecting the novel stimulus decreased as the number of trials after a novel stimulus was introduced increased (Monkey: *b*=-0.007, *95%CI*=-0.009 to -0.004, *p*<0.001; Human: *b*=-0.024, *95%CI*=-0.03 to -0.02, *p*<0.001; **Figure 1D-E**), whereas the probability of selecting the best alternative increased (Monkey: *b*=0.004, *95%CI*=0.002 to 0.006, *p*=0.001; Human: *b*=0.015, *95%*CI=0.01 to 0.02, *p*<0.001; **Figure 1D-E**). Collectively, behavioral performance indicated two common patterns across primate species: i) a novelty-seeking bias when making explore-exploit decisions, and ii) in time, this novelty bias wanes and primates learn to exploit the option with the highest assigned value.

**Figure 1.**
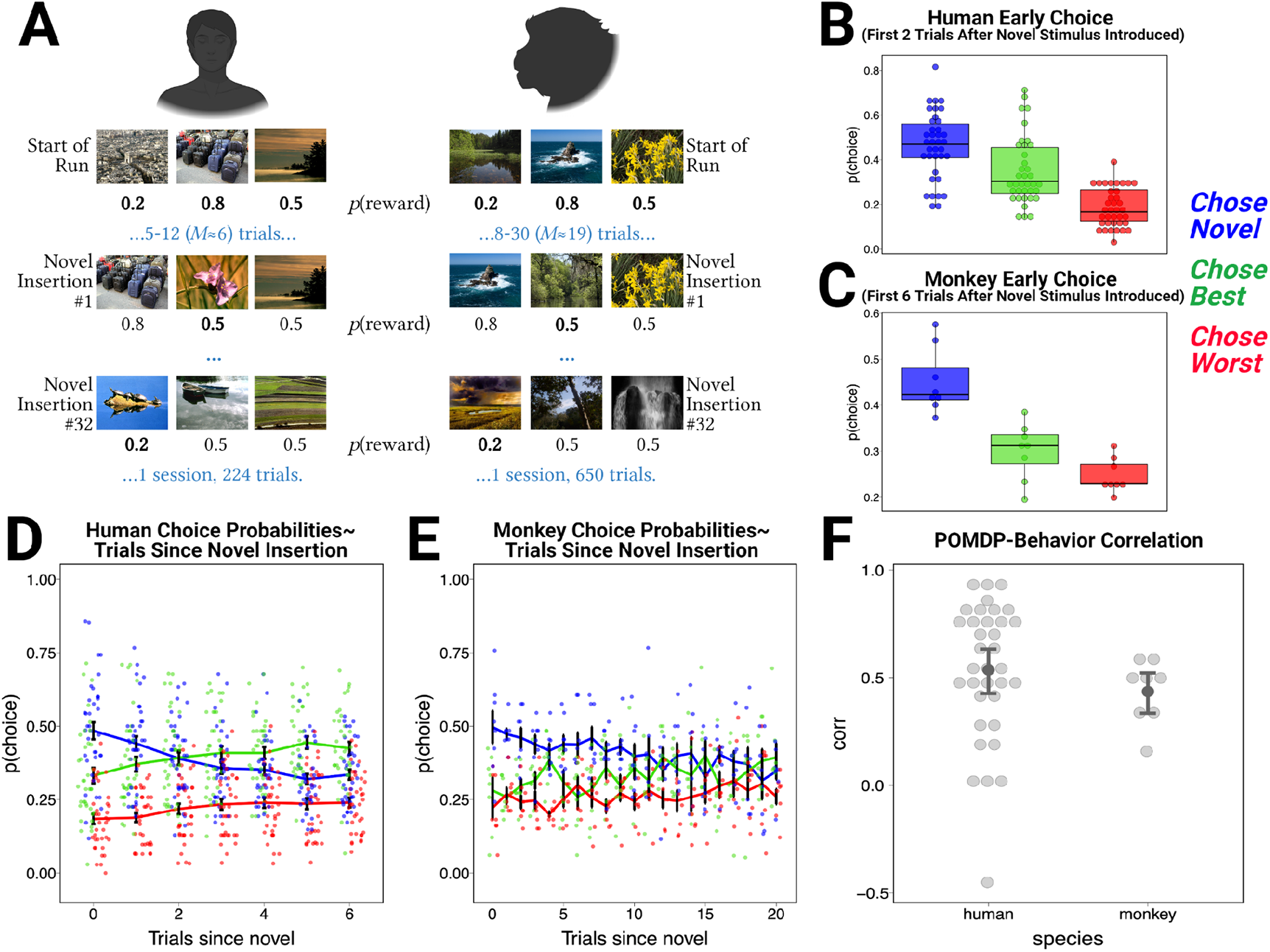
**(A)** Details of the novelty seeking three-arm bandit task session, as performed by human and monkey. Both humans and macaques chose between neutral images assigned the same nominal reward probabilities, and experienced the same number of overall novel stimulus insertion trials. Insertion rate was faster in humans. **(B-C)** Across both species, during the first few trials after a novel stimulus was introduced, selection of the novel stimulus increased relative to selection of the best or worst available alternative choice options. When they chose not to explore, both species also exploited the best alternative more often than choosing the worst available option. **(D-E)** Both humans and monkeys decreased their sampling of the novel option over time as a function of the number of trials elapsed since the insertion of a novel stimulus, and conversely both species also increased their selection of the best available option over time. **(F)** The correlation between behavioral task performance and the POMDP was significantly greater than zero within humans and monkeys, and POMDP-Behavior correlation strength did not differ between humans and monkeys, suggesting similar computations shape explore-exploit behavior across primate species.

We modeled normative explore-exploit policies as a Partially Observable Markov Decision Process (POMDP). In the POMDP, each option’s value is the sum of its immediate expected value (IEV) and future expected value (FEV). IEV reflects the likelihood that a particular choice will be rewarded. FEV is latent and reflects the sum of potential rewards to be earned in the future. Trial-to-trial changes in novelty-seeking are thought to be primarily shaped by an exploration BONUS, the relative difference in the FEV of an individual option relative to the FEV of all options. The exploration BONUS is highest when a novel option is first introduced, and decreases each time an option is sampled.

The POMDP value estimates were used in a non-linear regression approach to predict which option humans or monkeys would choose. In combination, the IEV and exploration BONUS associated with choices made by humans (*M*=0.59, *95%CI*=0.47 to 0.70, *t*=10.5, *p*<0.001) or rhesus macaques (*M*=0.45, *95%CI*=0.30 to 0.61, *t*=7.43, *p*<0.001) were predictive of overall performance. Examination of the individual regression coefficients in each species indicated that the IEV associated with each option was a strong determinant of whether or not an option was chosen (Human: *M*=0.44, *95%CI*=0.26 to 0.63, *t*=4.96, *p*<0.001; Monkey: *M*=0.12, *95%CI*=0.01 to 0.22, *t*=2.82, *p*=0.037). In humans, the exploration BONUS was not as strong as a predictor as it was in monkeys (Human: *M*=0.02, *95%CI*=-0.12 to 0.16, *t*=0.26, *p*=0.80; Monkey: *M*=0.08, *95%CI*=-0.002 to 0.16, *t*=2.52, *p*=0.05), perhaps because novelty was more salient to the monkeys due to differences in the number of trials that elapsed between the introduction of novel choice opportunities. However, in both humans and monkeys, we observed a negative correlation between IEV and BONUS regression coefficients (Human: *rho*=-0.48, *95%HDI*=-0.73 to -0.19; Monkey: *rho*=-0.67, *95%HDI*=-0.97 to -0.04). The strength of the POMDP-Behavior correlation did not differ across primate species (*M*_diff_=0.13, *95%CI*=-0.03 to 0.30, *t*=1.64, *p*=0.09; **Figure 1F**).

To elucidate the neurocomputational architecture of explore-exploit decision making in humans, we collected fMRI data and modeled choice-evoked responses as a function of value estimates, IEV and BONUS, derived from the POMDP model (Averbeck, 2015; Costa et al., 2019a). Bayesian multi-level modeling (Chen et al., 2019) was used to identify cortical and subcortical regions that encoded trial-by-trial changes in exploration BONUS and IEV associated with participants’ choices —given their importance in shaping exploration (Wittmann et al., 2008) and exploitation (Hunt et al., 2012), respectively. Several cortical and subcortical regions-of-interest (ROIs) encoded both relative IEV and BONUS, suggesting a role for these regions in shaping both exploitation and novelty-driven exploration. Both relative IEV and BONUS were associated with enhanced activation of medial FPC (mFPC), lateral OFC, subgenual anterior cingulate cortex (sgACC; **Figure 2**), and ventral subcortical regions (accumbens and amygdala; **Figure 3**). Conversely, both relative IEV and BONUS were associated with reduced activation of lateral FPC (lFPC; **Figure 2**).

**Figure 2.**
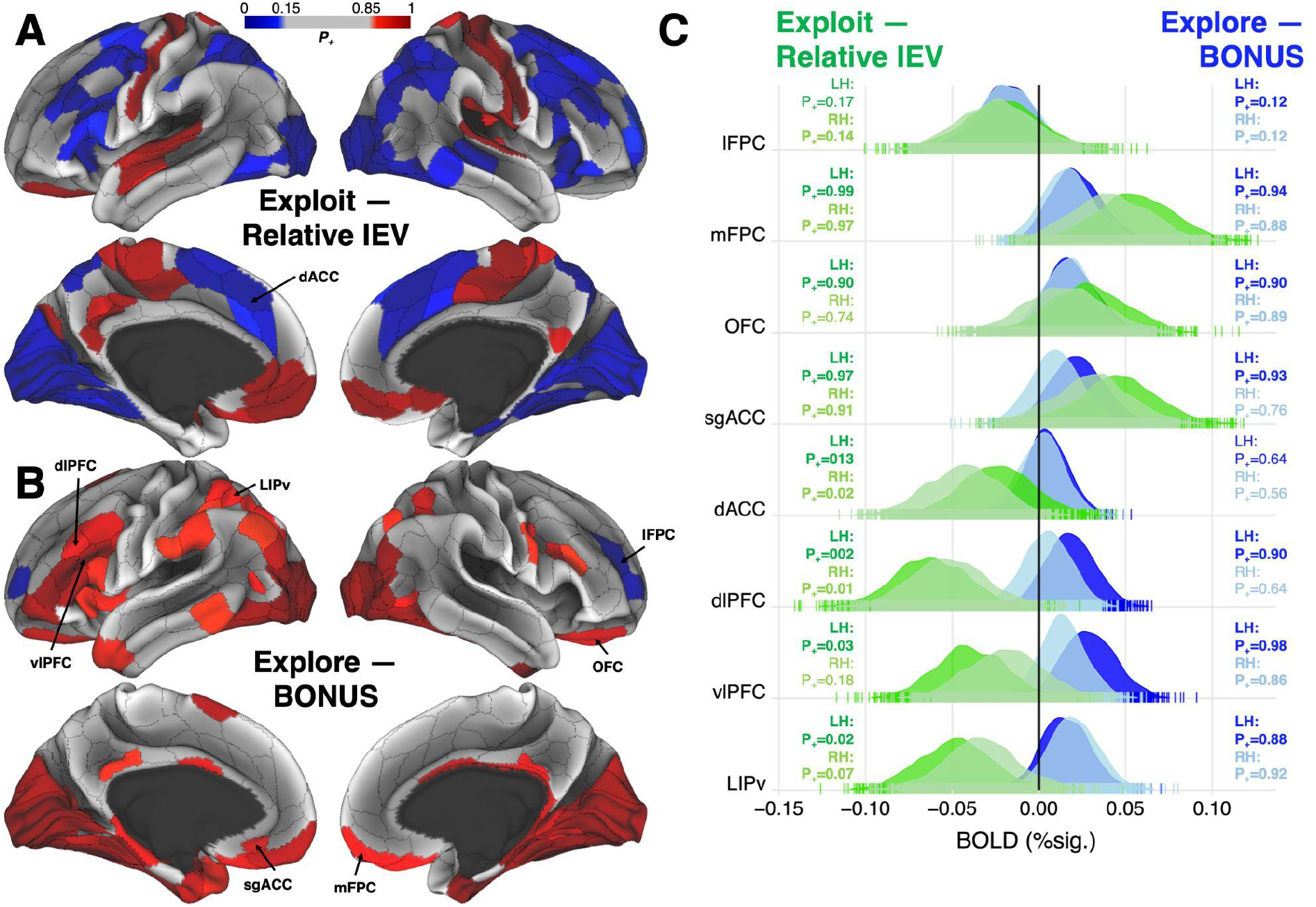
**(A)** Cortical regions showing credible evidence for positive (Red; ≥85% samples above 0) and negative (Blue; ≤15% samples above 0) encoding of an exploit-related computation (i.e., relative immediate expected value, IEV) **(B)** and an exploration / novelty-seeking computation (i.e., BONUS) in the Bayesian multilevel model. **(C)** Posterior distributions from subset of *a priori* regions-of-interest indicated a dissociation in frontopolar cortex between lateral FPC, which negatively encoded both relative IEV and BONUS suggesting reduced activation during both exploration and exploitation, and mFPC, OFC, and sgACC which positively encoded both parameters suggested *enhanced* activation during exploration and exploitation. Frontoparietal regions (namely: dACC, dlPFC, vlPFC, and LIPv) demonstrated within-region dissociations across exploit- and explore-related computations: Positive encoding of BONUS and negative encoding of relative IEV. Darker and lighter colors in posterior distributions indicate left- and right-hemispheres, respectively. Bolded text indicates either ≥85% or ≤15% of posterior samples above 0.

**Figure 3.**
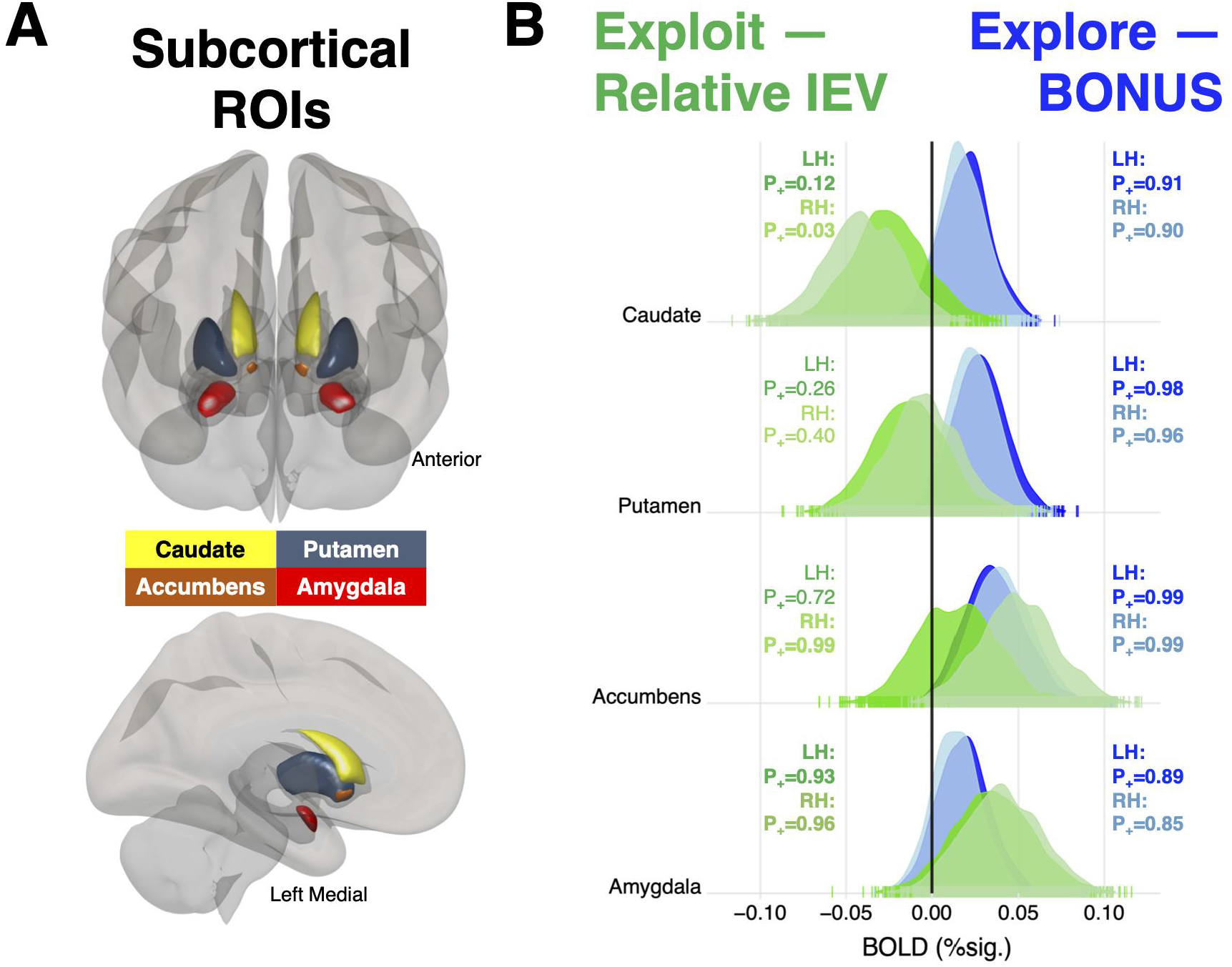
**(A)** Subset of the subcortical ROIs included in the Bayesian multilevel model. **(B)** Credible (≥85% samples above 0) evidence for positive encoding of relative IEV in accumbens and amygdala, while caudate negatively encoded relative IEV (≤15% samples above 0). In contrast, BONUS was positively encoded in dorsal striatum, accumbens, and amygdala. Darker and lighter colors in posterior distributions indicate left- and right-hemispheres, respectively. Bolded text indicates either ≥85% or ≤15% of posterior samples above 0.

In contrast, several regions demonstrated dissociable encoding of exploit- and explore-related computations. Specifically, whereas frontoparietal network regions (namely, dorsolateral prefrontal cortex, dlPFC; ventrolateral prefrontal cortex, vlPFC; area lateral intraparietal ventral, LIPv; and dorsal anterior cingulate cortex, dACC) demonstrated positive encoding of exploration BONUS, they negatively scaled with relative IEV (**Figure 2**). Dorsal striatal nuclei (caudate and putamen) positively encoded BONUS, and caudate negatively scaled with relative IEV (**Figure 3**). Sensorimotor responses also diverged across exploit- and explore-related computations: Visual areas showed enhanced activation as a function of BONUS and reduced activation as a function of relative IEV, whereas primary somatomotor cortices demonstrated enhanced activation as a function of relative IEV.

## Discussion

The current results provide key insight into the neural computations underlying explore-exploit decision making in primates. Similar to recent monkey neurophysiological data (Costa and Averbeck, 2020b; Costa et al., 2019a), we found encoding of explore- and exploit-related computations in regions known to be essential for reward-guided decision making—namely, ventral striatum [cf., (Wittmann et al., 2008)], amygdala, and ventromedial prefrontal regions. This suggests that homologous cortico-subcortical motivational neural circuits help to drive both reward-guided and novelty-seeking behaviors across primate species. Importantly, evidence for common neural computations shaping novelty-driven exploration across monkeys and humans should inform the development of next generation treatments for mental disorders associated with pathological explore-exploit behavior.

Encoding of POMDP derived valuations related to decisions to explore or exploit in lFPC indicated increased recruitment of lateral frontopolar cortex when exploration was motivated by a search for better choice opportunities rather than novelty seeking. Greater deactivation of lPFC occurred when participants exploited choice options they had repeatedly sampled and that they were certain would be rewarded at a high rate. But similar deactivation of lPFC was also observed when participants explored novel options whose reward value was uncertain. This implies that the lateral frontopolar cortex is deactivated when established policies surrounding decisions to exploit or explore are engaged. Moreover, using a POMDP to quantify the relative IEV and exploration bonuses associated with choosing a specific option, we found that deactivation of lPFC was inversely correlated with the relative immediate expected value and exploration bonuses of the monkeys’ choices. Taken together these results allow us to predict when lPFC is active in the context of explore-exploit decision making. Increased activation in lPFC should occur when participants chose relatively lower value options that had not yet been sampled or sampled infrequently several trials after a novel stimulus was last inserted (i.e., choices associated with a smaller exploration BONUS *and* low-to-moderate relative IEV). In human fMRI studies, lFPC is known to be activated in tasks that require multi-step planning or the control of responses according to competing goals (Braver et al., 2003; Koechlin et al., 1999). Negative encoding of BONUS and relative IEV at the time of choice in lFPC may indicate that this region plays a role in outcome monitoring and retaining the value of *unchosen* options (Boorman et al., 2009). An lFPC mechanism for outcome monitoring and encoding the value of unselected options at the time of choice could be critical for enabling primates’ ability to defer goals to explore new alternatives (Tsujimoto et al., 2011).

Lastly, we observed dissociations between explore-exploit computations in dorsal corticostriatal brain networks. Specifically, in the frontoparietal network and dorsal striatal nuclei (caudate and putamen), BOLD activity was increased as a function of BONUS, suggesting a role in computing the latent value of exploring novel stimuli. This function would complement the already known role of frontoparietal regions in matching behavior (Sugrue et al., 2004), foraging (Genovesio, Wise, and Passingham, 2014), and information sampling (Furl and Averbeck, 2011; Costa and Averbeck, 2015). Several of these frontoparietal and dorsal striatal regions were also *negatively* associated with relative IEV, indicating deactivation during the exploitation of familiar rewards. One possibility is that participants rapidly form an internal model of the general structure of the task, which includes an expectation about the substitution rate that controls how often a novel choice option is introduced. This acquired model of the task would enable the subject to infer that novel stimuli provide a potential increase in future value relative to familiar options (in accordance with the BONUS parameter from the POMDP). Conversely, after participants have had the opportunity to repeatedly sample novel choice options and learn whether they are better or worse than familiar alternatives (i.e., when all options are familiar, and their relative IEV is differentiated), participants may begin making decisions using a model-free stimulus-outcome learning system. Therefore, positive encoding of BONUS and negative encoding of relative IEV in dorsal frontoparietal and dorsal striatal regions would accord with the view that these regions play a role in Bayesian state inference and model-based explore-exploit decision making (Bartolo and Averbeck, 2020). Collectively, these findings should motivate future single-cell recording studies in lFPC, frontoparietal circuits, and dorsal striatum during explore-exploit behavior in nonhuman primates.

Overall, the current study provides compelling evidence that humans and monkeys perform similar neural computations when exploring novel stimuli in lieu of exploiting familiar rewards. Specifically, across primate species we observed a novelty-seeking decision bias in the immediate wake of encountering new choice opportunities, alongside an increased tendency to exploit the best available option relative to the worst available alternative. These decision tendencies were well-modeled as a POMDP across both human participants and monkey subjects, with the model fit not differing significantly across species. Therefore, novelty-seeking under explore-exploit tensions represents an exciting avenue for cross-species primate research, providing a new bench-to-bedside pipeline for interventions for pathological reward processing or novelty sensitivity in clinical populations (e.g. substance use disorder).

## Methods

### Human Participants

47 adult participants from the Albuquerque, New Mexico community were enrolled in the current study (**Supplementary Table 1**), which was approved through the University of New Mexico Office of the Institutional Review Board. Unfortunately, *N*=5 of 47 enrolled participants were unable to complete their scheduled study visit due to mandated COVID19 pandemic lockdown restrictions in Spring, 2020. Additionally, *N*=1 participant was removed from study analyses due to extreme non-normative behavior relative to the computational model (3.34 residual standard deviations from the group line of best fit between the BONUS and IEV coefficients), *N*=1 participant was removed due to excessive head motion during fMRI (>3 standard deviations above mean framewise displacement across all task fMRI runs), *N*=2 participants were removed due to insufficient responding during the task (*N*=144 responses during the task, -4.11 standard deviations below mean across the sample), and *N*=1 participant was removed due to an incorrect key configuration on the task fMRI button box. Therefore, the final sample in the current study comprised *N*=37 participants (**Supplementary Table 2**).

### Monkey Subjects

Eight adult male rhesus macaques (*Macca mulatta*) served as subjects. Their ages and weights at the start of training ranged between 6-8 years and 7.2-9.3 kg. Animals were pair housed when possible, had access to food 24 hours a day, were kept on a 12-h light-dark cycle, and tested during the light portion of the day. On testing days, the monkeys earned their fluid through performance on the task, whereas on non-testing days the animals were given free access to water. All procedures were reviewed and approved by the NIMH Animal Care and Use Committee. Behavioral data from a subset of the same monkeys also appears in Costa et al. 2019 (Costa et al., 2019b).

### Novelty-Bandit Task

#### Human version

Participants made speeded (≤2 seconds) manual responses between three neutral images taken from the International Affective Pictures System (IAPS(Bradley and Lang, 2020)). Stimuli were presented using EPrime Version 3 (Psychology Software Tools, Sharpsburg, PA), with responses recorded using a MIND Input Device (https://www.mrn.org/collaborate/mind-input-device). Images were randomly assigned an *a priori* low (*p*=0.2), medium (*p*=0.5), or high (*p*=0.8) reward probability. Every 5-12 trials (*M*≈6) a ‘novel insertion’ took place, wherein one familiar image in the current set was replaced by one novel image to create a new set that would be presented for the proceeding 5-12 trials. Novel images were randomly assigned a low, medium, or high reward probability, with the caveat that all 3 images could not have the same assigned reward probability in the new set. Participants completed 224 trials containing 32 novel stimulus insertions, divided evenly into 4*≈7-minute fMRI runs. Participants made confidence judgments (low, medium, or high confidence) after each decision, but these data are not relevant to the current manuscript. Image location was randomized on each trial, and participants received either reward (green ‘+1’) or nonreward (red ‘0’) feedback after each decision.

#### Monkey version

Subjects performed a saccade-based version of the same task performed by humans. Stimulus presentation and behavioral assessment were controlled via Monkeylogic (Hwang et al., 2019), and eye movements were sampled at 400 frames per second, 1000 Hz using an Arrington Viewpoint eye tracker (Arrington Research, Scottsdale, AZ). Each session began with three naturalistic scenes randomly assigned an *a priori* low, medium, or high probability of being paired with an apple juice reward via a precise liquid-delivery device (Mitz, 2005). Every 8-30 trials (*M*≈19 trials) a novel insertion took place, and the novel image was assigned a low, medium, or high reward probability, with the caveat that all three images in the new set could not have the same reward likelihood. Monkeys completed 650 trials per session, with each trial beginning with a central fixation (250-750ms), followed by the presentation of three images in the periphery, and a saccade to and fixation on one of the targets for 500ms). Subjects performed a saccade-based version of the same task performed by humans. Stimulus presentation and behavioral assessment were controlled via Monkeylogic (Hwang et al., 2019), and eye movements were sampled at 400 frames per second, 1000 Hz using an Arrington Viewpoint eye tracker (Arrington Research, Scottsdale, AZ). Each session began with three naturalistic scenes randomly assigned an *a priori* low, medium, or high probability of being paired with an apple juice reward via a precise liquid-delivery device (Mitz, 2005). Every 8-30 trials (*M*≈19 trials) a novel insertion took place, and the novel image was assigned a low, medium, or high reward probability, with the caveat that all three images in the new set could not have the same reward likelihood. Monkeys completed 650 trials per session, with each trial beginning with a central fixation (250-750ms), followed by the presentation of three images in the periphery, and a saccade to and fixation on one of the targets for 500ms).

### Magnetic Resonance Imaging (MRI) Acquisition, Processing, and Analysis

#### Image acquisition

All MRI scans were acquired on a 3T Siemens Tim Trio system with a 32-channel phased-array head coil. T1-weighted (T1w) structural MRI was acquired via a multi-echo MPRAGE sequence (5-echo; voxel size=1mm iso). T2*-weighted functional MRI data were acquired with a gradient EPI pulse sequence using simultaneous multi-slice technology (TR=1s; TE=30ms; Flip=44°; MB factor=4; voxel size=3mm iso). Acquired data were converted from DICOM to Brain Imaging Data Structure (BIDS) format([CSL STYLE ERROR: reference with no printed form.]) using Heudiconv v.0.5.4 (https://heudiconv.readthedocs.io/en/latest/).

#### Image preprocessing

Results included in this manuscript come from preprocessing performed using fMRIPrep 20.2.0rc0 (Esteban et al., 2019) (RRID:SCR_016216), which is based on Nipype 1.5.1 (Gorgolewski et al., 2011) (RRID:SCR_002502). Full preprocessing pipeline details can be found in **Supplementary Methods**.

#### fMRI analysis

##### First-level model

Participant-level fMRI data were modeled as a function of parameters from the Partially Observable Markov Decision Process (POMDP) model. Specifically, in a first pass model we fit the fMRI timeseries as a function of six regressors: 1) exploration bonus (BONUS), 2) immediate expected value (IEV), 3) future expected value (FEV), 4) number of trials since a novel stimulus was inserted, 5) whether or not the previous trial was rewarded, and 6) reward prediction error (feedback 1 or 0, minus IEV). Additionally, given the convention of the field to look at *relative* IEV as an indicator of the neural circuits underlying value-based choice (Chau et al., 2014; Hunt et al., 2012), we ran a second model replacing the IEV regressor with a relIEV parameter reflecting the difference between the chosen option’s IEV and the average IEV of the two alternatives. In both models, regressors 1-6 were linear optimal basis sets time-locked to the choice event, whereas regressor 6 was locked to the feedback event. Notably, multicollinearity was not a concern in either model (all VIFs≤1.54). Cue onset timing was jittered and optimized via AFNI’s make_random_timings.py. Here, we report the full results from second-level models for all POMDP regressors (BONUS, Relative IEV, IEV, FEV, and RPE) for the same subset of ROIs focused on in the main text (**Supplementary Tables 3-7**). 24 standard head motion parameters and their derivatives were included as confound regressors(Satterthwaite et al., 2013), and 5mm full-width half-maximum smoothing and 100s high-pass temporal filtering were applied to the first-level data.

##### Second-level model

Second-level data were modeled using a Bayesian Multilevel Modelling approach using brms v2.15.0 (https://cran.r-project.org/web/packages/brms/index.html) in R v4.0.4 (https://www.r-project.org). Relative to conventional mass univariate analyses, Bayesian Multilevel Modelling improves model efficiency and sensitivity for detecting effects at smaller brain regions(Chen et al., 2019, [CSL STYLE ERROR: reference with no printed form.]). First, mean percent signal change was extracted for each participant from 370 anatomically-defined regions-of-interest (ROIs) spanning cortex and subcortex. Specifically, cortex was parcellated using 360 areas (180 from each hemisphere) from the Glasser Multimodal Parcellation (Glasser et al., 2016), and 10 subcortical ROIs comprising amygdala and basal ganglia were segmented using the Harvard-Oxford Probabilistic Atlas via the FMRIB Software Library (FSL; (Smith et al., 2001, 2004)). Next, second-level models were fit in brms for each first-level model parameter of interest in the form:

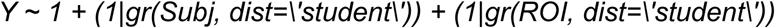

In these models, *Y* corresponds to the percent signal change to a given regressor from the first-level models, the *Subj* term represents the random effect associated with each subject, and the *ROI* term represents the random effect associated with each cortical and subcortical ROI in the model. All brms models were re-run incorporating measurement errors for each *Y*-value, but this did not alter the inferences described in the main text. Notably, we used non-informative priors.

Lastly, we extracted the marginal posteriors associated with each ROI. Importantly, the main output of each brms model is one overall posterior distribution that is a joint distribution across participants and ROIs in a high-dimensional parameter space, and therefore correction for multiple comparisons is not appropriate (Gelman et al., 2012; Limbachia et al., 2021). Statistical inferences regarding the credibility of each ROI encoding a given regressor were made based on the proportion of each distribution that was above 0 (henceforth, *P+*). *P+* values less than 0.15 were used to indicate credible evidence for negative encoding of a given regressor, whereas *P+* values above 0.85 indicated credible evidence for positive encoding. Though “strong,” “moderate,” and “weak” labels are sometimes assigned to arbitrary *P+* levels within a Bayesian multilevel modelling framework, our goal in the current study was to map the brainwide computational architecture of explore-exploit decision making for the first time in humans. Therefore, even ROIs demonstrating “weak” positive or negative encoding of a given regressor could be theoretically important, and therefore worthy of discussion in the main text (Limbachia et al., 2021).

## Footnotes

1 ”Early trials” were defined as the first N=2 trials after a novel stimulus was introduced for humans. Relative to the typical interval between insertion trials (M≈6 for humans, M≈19 for monkeys), this was translated to N=6 trials for the monkeys.

