## Supplementary Materials for "The neurocomputational architecture of explore-exploit decision making"

**Title:** SUPPLEMENTARY MATERIALS: The neurocomputational architecture of explore-exploit decision making in humans.

**Affiliations:**

### Supplementary Methods

#### Image Preprocessing

Results included in this manuscript come from preprocessing performed using fMRIPrep 20.2.0rc0 (Esteban et al., 2019) (RRID:SCR_016216), which is based on Nipype 1.5.1 (Gorgolewski et al., 2011) (RRID:SCR_002502).

**Anatomical data preprocessing**. The T1w image was corrected for intensity non-uniformity (INU) with N4BiasFieldCorrection (Tustison et al., 2010), distributed with ANTs 2.2.0 (Avants et al., 2008) (RRID:SCR_004757), and used as T1w-reference throughout the workflow. The T1w-reference was then skull-stripped with a Nipype implementation of the antsBrainExtraction.sh workflow (from ANTs), using OASIS30ANTs as target template. Brain tissue segmentation of cerebrospinal fluid (CSF), white-matter (WM) and gray-matter (GM) was performed on the brain-extracted T1w using fast (FSL 5.0.9; (Avants et al., 2008; Zhang et al., 2001), RRID:SCR_002823). Brain surfaces were reconstructed using recon-all (FreeSurfer 6.0.1; (Dale et al., 1999), RRID:SCR_001847), and the brain mask estimated previously was refined with a custom variation of the method to reconcile ANTs-derived and FreeSurfer-derived segmentations of the cortical gray-matter of Mindboggle (Dale et al., 1999; Klein et al., 2017) (RRID:SCR_002438). Volume-based spatial normalization to one standard space (MNI152NLin2009cAsym) was performed through nonlinear registration with antsRegistration (ANTs 2.2.0), using brain-extracted versions of both T1w reference and the T1w template. The following template was selected for spatial normalization: ICBM 152 Nonlinear Asymmetrical template version 2009c (Fonov et al., 2009) [RRID:SCR_008796; TemplateFlow ID: MNI152NLin2009cAsym],

**Functional data preprocessing**. First, a reference volume and its skull-stripped version were generated using a custom methodology of fMRIPrep. A deformation field to correct for susceptibility distortions was estimated based on fMRIPrep's fieldmap-less approach. The deformation field is that resulting from co-registering the BOLD reference to the same-subject T1w-reference with its intensity inverted (Wang et al., 2017). Registration is performed with antsRegistration (ANTs 2.2.0), and the process regularized by constraining deformation to be nonzero only along the phase-encoding direction, and modulated with an average fieldmap template (Treiber et al., 2016). Based on the estimated susceptibility distortion, a corrected EPI (echo-planar imaging) reference was calculated for a more accurate co-registration with the anatomical reference. The BOLD reference was then co-registered to the T1w reference using bbregister (FreeSurfer) which implements boundary-based registration (Greve and Fischl, 2009). Co-registration was configured with six degrees of freedom. Head-motion parameters with respect to the BOLD reference (transformation matrices, and six corresponding rotation and translation parameters) are estimated before any spatiotemporal filtering using mcflirt (FSL 5.0.9; (Jenkinson et al., 2002)). BOLD runs were slice-time corrected using 3dTshift from AFNI 20160207; (Cox and Hyde, 1997) (RRID:SCR_005927). The BOLD time-series (including slice-timing correction when applied) were resampled onto their original, native space by applying a single, composite transform to correct for head-motion and susceptibility distortions. These resampled BOLD time-series will be referred to as preprocessed BOLD in original space, or just preprocessed BOLD. The BOLD time-series were resampled into standard space, generating a preprocessed BOLD run in MNI152NLin2009cAsym space. First, a reference volume and its skull-stripped version were generated using a custom methodology of fMRIPrep. Several confounding time-series were calculated based on the preprocessed BOLD: framewise displacement (FD), DVARS and three region-wise global signals. FD was computed using two formulations following Power (absolute sum of relative motions; (Power et al., 2014)) and

Jenkinson (relative root mean square displacement between affines; (Jenkinson et al., 2002)). FD and DVARS are calculated for each functional run, both using their implementations in Nipype (following the definitions by Power et al. 2014). The three global signals are extracted within the CSF, the WM, and the whole-brain masks. Additionally, a set of physiological regressors were extracted to allow for component-based noise correction (CompCor; (Behzadi et al., 2007)). Principal components are estimated after high-pass filtering the preprocessed BOLD time-series (using a discrete cosine filter with 128s cut-off) for the two CompCor variants: temporal (tCompCor) and anatomical (aCompCor). tCompCor components are then calculated from the top 2% variable voxels within the brain mask. For aCompCor, three probabilistic masks (CSF, WM and combined CSF+WM) are generated in anatomical space. The implementation differs from that of Behzadi et al. in that instead of eroding the masks by 2 pixels on BOLD space, the aCompCor masks are subtracted from a mask of pixels that likely contain a volume fraction of GM. This mask is obtained by dilating a GM mask extracted from the FreeSurfer's aseg segmentation, and it ensures components are not extracted from voxels containing a minimal fraction of GM. Finally, these masks are resampled into BOLD space and binarized by thresholding at 0.99 (as in the original implementation). Components are also calculated separately within the WM and CSF masks. For each CompCor decomposition, the k components with the largest singular values are retained, such that the retained components' time series are sufficient to explain 50 percent of variance across the nuisance mask (CSF, WM, combined, or temporal). The remaining components are dropped from consideration. The head-motion estimates calculated in the correction step were also placed within the corresponding confounds file. The confound time series derived from head motion estimates and global signals were expanded with the inclusion of temporal derivatives and quadratic terms for each (Satterthwaite et al., 2013). Frames that exceeded a threshold of 0.5 mm FD or 1.5 standardised DVARS were annotated as motion outliers. All resamplings can be performed with a single interpolation step by composing all the pertinent transformations (i.e. head-motion transform matrices, susceptibility distortion correction when available, and co-registrations to anatomical and output spaces). Gridded (volumetric) resamplings were performed using antsApplyTransforms (ANTs), configured with Lanczos interpolation to minimize the smoothing effects of other kernels (Lanczos, 1964). Non-gridded (surface) resamplings were performed using mri_vol2surf (FreeSurfer). Many internal operations of fMRIPrep use Nilearn 0.6.2 (Abraham et al., 2014) (RRID:SCR_001362), mostly within the functional processing workflow. For more details of the pipeline, see the section corresponding to workflows in fMRIPrep's documentation.

**Copyright Waiver**. The above boilerplate text was automatically generated by fMRIPrep with the express intention that users should copy and paste this text into their manuscripts unchanged. It is released under the CC0 license.

### Supplementary Tables

**Supplementary Table 1.** Participant demographics for all participants enrolled in the Human Subjects fMRI study.

| | |
|---|---|
| ***Sex*** | |
| *Female (N)* | 28 |
| *Male (N)* | 19 |
| ***Age*** | |
| *Mean±SD* | 28.0 ± 8.81 |
| ***Ethnicity*** | |
| *Hispanic or Latino (N)* | 20 |
| *Not Hispanic or Latino (N)* | 22 |
| *Unknown/Not Reported (N)* | 5 |
| ***Race*** | |
| *American Indian/Alaska Native, Black or African American (N)* | 1 |
| *American Indian/Alaska Native, White (N)* | 2 |
| *Asian (N)* | 4 |
| *Black or African American (N)* | 1 |
| *White (N)* | 29 |
| *Unknown/Not Reported (N)* | 10 |

| **Supplementary Table 2.** Participant demographics for all participants included in the final dataset for the Human Subjects study. | |
|---|---|
| ***Sex*** | |
| *Female (N)* | 24 |
| *Male (N)* | 13 |
| ***Age*** | |
| *Mean±SD* | 26.6 ± 7.24 |
| ***Ethnicity*** | |
| *Hispanic or Latino (N)* | 12 |
| *Not Hispanic or Latino (N)* | 20 |
| *Unknown/Not Reported (N)* | 5 |
| ***Race*** | |
| *American Indian/Alaska Native, Black or African American (N)* | 1 |
| *American Indian/Alaska Native, White (N)* | 1 |
| *Asian (N)* | 3 |
| *Black or African American (N)* | 1 |
| *White (N)* | 26 |
| *Unknown/Not Reported (N)* | 5 |

**Supplementary Table 3.** Summary table for exploration bonus (BONUS) Bayesian Multilevel Model. Blue indicates regions with *P+* ≤ 0.15 and red indicates *P+* ≥ 0.85.

| ROI | *Mean* | *SE* | *P+* | *0.25* | *0.05* | *0.50* | *0.95* | *0.975* |
|---|---|---|---|---|---|---|---|---|
| ***Cortical Regions*** | | | | | | | | |
| *LH_p10p* | -0.0192 | 0.0004 | 0.1155 | -0.0529 | -0.0478 | -0.0188 | 0.0071 | 0.0117 |
| *RH_p10p* | -0.0186 | 0.0004 | 0.1255 | -0.0507 | -0.0450 | -0.0185 | 0.0074 | 0.0118 |
| *LH_pOFC* | -0.0065 | 0.0004 | 0.3465 | -0.0404 | -0.0342 | -0.0060 | 0.0180 | 0.0219 |
| *RH_4* | 0.0008 | 0.0003 | 0.5325 | -0.0249 | -0.0206 | 0.0011 | 0.0208 | 0.0258 |
| *RH_a32pr* | 0.0016 | 0.0003 | 0.5620 | -0.0251 | -0.0205 | 0.0021 | 0.0230 | 0.0265 |
| *RH_pOFC* | 0.0021 | 0.0003 | 0.5715 | -0.0304 | -0.0249 | 0.0025 | 0.0265 | 0.0317 |
| *LH_4* | 0.0036 | 0.0003 | 0.6255 | -0.0211 | -0.0178 | 0.0037 | 0.0236 | 0.0282 |
| *RH_p9-46v* | 0.0049 | 0.0003 | 0.6430 | -0.0221 | -0.0182 | 0.0051 | 0.0276 | 0.0313 |
| *LH_a32pr* | 0.0043 | 0.0003 | 0.6440 | -0.0211 | -0.0167 | 0.0041 | 0.0251 | 0.0295 |
| *RH_s32* | 0.0098 | 0.0003 | 0.7575 | -0.0181 | -0.0137 | 0.0098 | 0.0337 | 0.0390 |
| *LH_V4t* | 0.0120 | 0.0003 | 0.8050 | -0.0163 | -0.0110 | 0.0117 | 0.0355 | 0.0415 |
| *RH_IFSp* | 0.0137 | 0.0003 | 0.8555 | -0.0130 | -0.0077 | 0.0135 | 0.0356 | 0.0398 |
| *LH_LIPv* | 0.0156 | 0.0003 | 0.8785 | -0.0121 | -0.0073 | 0.0150 | 0.0389 | 0.0441 |
| *RH_10v* | 0.0171 | 0.0003 | 0.8835 | -0.0110 | -0.0068 | 0.0167 | 0.0429 | 0.0471 |
| *RH_OFC* | 0.0176 | 0.0003 | 0.8940 | -0.0100 | -0.0053 | 0.0176 | 0.0419 | 0.0477 |
| *LH_p9-46v* | 0.0182 | 0.0003 | 0.9030 | -0.0088 | -0.0044 | 0.0180 | 0.0415 | 0.0461 |

| | | | | | | | | |
|---|---|---|---|---|---|---|---|---|
| *LH_OFC* | 0.0185 | 0.0003 | 0.9050 | -0.0093 | -0.0048 | 0.0178 | 0.0429 | 0.0478 |
| *RH_LIPv* | 0.0202 | 0.0003 | 0.9215 | -0.0083 | -0.0032 | 0.0196 | 0.0444 | 0.0497 |
| *LH_s32* | 0.0228 | 0.0004 | 0.9310 | -0.0060 | -0.0019 | 0.0224 | 0.0493 | 0.0556 |
| *LH_10v* | 0.0220 | 0.0003 | 0.9445 | -0.0045 | -0.0005 | 0.0207 | 0.0480 | 0.0547 |
| *LH_IFSp* | 0.0292 | 0.0003 | 0.9760 | 0.0007 | 0.0050 | 0.0287 | 0.0553 | 0.0606 |
| *RH_V4t* | 0.0346 | 0.0004 | 0.9855 | 0.0025 | 0.0070 | 0.0331 | 0.0662 | 0.0714 |
| *RH_V4* | 0.0604 | 0.0006 | 0.9950 | 0.0127 | 0.0184 | 0.0591 | 0.1053 | 0.1123 |
| *LH_V4* | 0.0465 | 0.0005 | 0.9955 | 0.0092 | 0.0149 | 0.0444 | 0.0840 | 0.0905 |
| ***Subcortical Regions*** | | | | | | | | |
| *RH_Pallidum* | 0.0073 | 0.0003 | 0.7185 | -0.0191 | -0.0151 | 0.0071 | 0.0302 | 0.0342 |
| *RH_Amygdala* | 0.0139 | 0.0003 | 0.8470 | -0.0140 | -0.0091 | 0.0138 | 0.0383 | 0.0415 |
| *LH_Pallidum* | 0.0133 | 0.0003 | 0.8565 | -0.0133 | -0.0078 | 0.0131 | 0.0342 | 0.0383 |
| *LH_Amygdala* | 0.0182 | 0.0003 | 0.8865 | -0.0109 | -0.0058 | 0.0180 | 0.0446 | 0.0500 |
| *RH_Caudate* | 0.0174 | 0.0003 | 0.9000 | -0.0093 | -0.0047 | 0.0166 | 0.0411 | 0.0456 |
| *LH_Caudate* | 0.0193 | 0.0003 | 0.9070 | -0.0085 | -0.0042 | 0.0198 | 0.0442 | 0.0485 |
| *RH_Putamen* | 0.0237 | 0.0003 | 0.9585 | -0.0028 | 0.0012 | 0.0233 | 0.0467 | 0.0514 |
| *LH_Putamen* | 0.0280 | 0.0003 | 0.9780 | 0.0004 | 0.0035 | 0.0275 | 0.0532 | 0.0584 |
| *LH_Accumbens* | 0.0364 | 0.0004 | 0.9875 | 0.0035 | 0.0089 | 0.0351 | 0.0672 | 0.0755 |
| *RH_Accumbens* | 0.0404 | 0.0004 | 0.9935 | 0.0084 | 0.0131 | 0.0397 | 0.0701 | 0.0758 |

**Supplementary Table 4.** Summary table for Relative Immediate Expected Value (Relative IEV) Bayesian Multilevel Model. Blue indicates regions with *P+* ≤ 0.15 and red indicates *P+* ≥ 0.85.

| ROI | *Mean* | *SE* | *P+* | *0.25* | *0.05* | *0.50* | *0.95* | *0.975* |
|---|---|---|---|---|---|---|---|---|
| ***Cortical Regions*** | | | | | | | | |
| *LH_p9-46v* | -0.0620 | 0.0005 | 0.0025 | -0.1049 | -0.0974 | -0.0624 | -0.0270 | -0.0168 |
| *RH_V4* | -0.0624 | 0.0005 | 0.0045 | -0.1098 | -0.1022 | -0.0619 | -0.0220 | -0.0161 |
| *RH_p9-46v* | -0.0520 | 0.0005 | 0.0100 | -0.0942 | -0.0874 | -0.0526 | -0.0143 | -0.0087 |
| *LH_LIPv* | -0.0476 | 0.0005 | 0.0205 | -0.0887 | -0.0831 | -0.0475 | -0.0110 | -0.0017 |
| *RH_a32pr* | -0.0433 | 0.0005 | 0.0240 | -0.0853 | -0.0785 | -0.0431 | -0.0091 | -0.0012 |
| *LH_V4* | -0.0473 | 0.0005 | 0.0245 | -0.0936 | -0.0858 | -0.0471 | -0.0089 | -0.0010 |
| *RH_V4t* | -0.0429 | 0.0005 | 0.0275 | -0.0853 | -0.0783 | -0.0429 | -0.0071 | 0.0004 |
| *LH_IFSp* | -0.0392 | 0.0005 | 0.0335 | -0.0806 | -0.0725 | -0.0397 | -0.0052 | 0.0034 |
| *RH_LIPv* | -0.0334 | 0.0005 | 0.0660 | -0.0764 | -0.0690 | -0.0334 | 0.0032 | 0.0086 |
| *LH_a32pr* | -0.0238 | 0.0005 | 0.1290 | -0.0652 | -0.0585 | -0.0239 | 0.0120 | 0.0185 |
| *RH_p10p* | -0.0237 | 0.0005 | 0.1430 | -0.0658 | -0.0596 | -0.0238 | 0.0142 | 0.0205 |
| *LH_p10p* | -0.0206 | 0.0005 | 0.1710 | -0.0637 | -0.0580 | -0.0208 | 0.0164 | 0.0242 |
| *RH_IFSp* | -0.0196 | 0.0005 | 0.1830 | -0.0613 | -0.0547 | -0.0193 | 0.0155 | 0.0216 |
| *LH_V4t* | -0.0057 | 0.0005 | 0.4025 | -0.0498 | -0.0414 | -0.0061 | 0.0322 | 0.0404 |
| *RH_OFC* | 0.0150 | 0.0005 | 0.7385 | -0.0301 | -0.0234 | 0.0148 | 0.0533 | 0.0606 |
| *RH_pOFC* | 0.0208 | 0.0006 | 0.8090 | -0.0300 | -0.0212 | 0.0210 | 0.0611 | 0.0691 |

| | | | | | | | | |
|---|---|---|---|---|---|---|---|---|
| *LH_OFC* | 0.0280 | 0.0005 | 0.8980 | -0.0164 | -0.0094 | 0.0277 | 0.0650 | 0.0714 |
| *RH_s32* | 0.0306 | 0.0005 | 0.9075 | -0.0143 | -0.0070 | 0.0312 | 0.0677 | 0.0753 |
| *LH_4* | 0.0270 | 0.0005 | 0.9085 | -0.0134 | -0.0078 | 0.0272 | 0.0607 | 0.0669 |
| *LH_s32* | 0.0450 | 0.0005 | 0.9670 | -0.0035 | 0.0055 | 0.0452 | 0.0826 | 0.0899 |
| *RH_4* | 0.0412 | 0.0005 | 0.9690 | -0.0021 | 0.0054 | 0.0416 | 0.0767 | 0.0837 |
| *RH_10v* | 0.0418 | 0.0005 | 0.9740 | -0.0001 | 0.0049 | 0.0415 | 0.0791 | 0.0881 |
| *LH_10v* | 0.0549 | 0.0005 | 0.9920 | 0.0097 | 0.0178 | 0.0547 | 0.0925 | 0.0983 |
| *LH_pOFC* | 0.0646 | 0.0006 | 0.9950 | 0.0164 | 0.0244 | 0.0641 | 0.1071 | 0.1138 |
| ***Subcortical Regions*** | | | | | | | | |
| *RH_Caudate* | -0.0400 | 0.0005 | 0.0305 | -0.0834 | -0.0766 | -0.0402 | -0.0043 | 0.0023 |
| *LH_Caudate* | -0.0251 | 0.0005 | 0.1240 | -0.0670 | -0.0602 | -0.0255 | 0.0112 | 0.0185 |
| *RH_Pallidum* | -0.0203 | 0.0005 | 0.1685 | -0.0626 | -0.0550 | -0.0207 | 0.0171 | 0.0222 |
| *LH_Pallidum* | -0.0129 | 0.0005 | 0.2535 | -0.0541 | -0.0478 | -0.0130 | 0.0221 | 0.0285 |
| *LH_Putamen* | -0.0135 | 0.0005 | 0.2585 | -0.0552 | -0.0480 | -0.0135 | 0.0207 | 0.0294 |
| *RH_Putamen* | -0.0047 | 0.0005 | 0.4025 | -0.0475 | -0.0388 | -0.0048 | 0.0300 | 0.0379 |
| *LH_Accumbens* | 0.0144 | 0.0005 | 0.7190 | -0.0306 | -0.0231 | 0.0143 | 0.0530 | 0.0623 |
| *LH_Amygdala* | 0.0353 | 0.0005 | 0.9300 | -0.0114 | -0.0031 | 0.0350 | 0.0747 | 0.0823 |
| *RH_Amygdala* | 0.0402 | 0.0005 | 0.9645 | -0.0035 | 0.0035 | 0.0400 | 0.0778 | 0.0852 |
| *RH_Accumbens* | 0.0499 | 0.0005 | 0.9870 | 0.0053 | 0.0124 | 0.0493 | 0.0883 | 0.0951 |

**Supplementary Table 5.** Summary table for Immediate Expected Value (IEV) Bayesian Multilevel Model. Blue indicates regions with *P+* ≤ 0.15 and red indicates *P+* ≥ 0.85.

| ROI | *Mean* | *SE* | *P+* | *0.25* | *0.05* | *0.50* | *0.95* | *0.975* |
|---|---|---|---|---|---|---|---|---|
| ***Cortical Regions*** | | | | | | | | |
| *LH_p9-46v* | -0.0337 | 0.0005 | 0.0620 | -0.0782 | -0.0696 | -0.0342 | 0.0028 | 0.0128 |
| *RH_p9-46v* | -0.0232 | 0.0005 | 0.1465 | -0.0665 | -0.0583 | -0.0231 | 0.0123 | 0.0212 |
| *RH_a32pr* | -0.0216 | 0.0005 | 0.1540 | -0.0624 | -0.0554 | -0.0221 | 0.0147 | 0.0218 |
| *LH_IFSp* | -0.0212 | 0.0005 | 0.1560 | -0.0630 | -0.0577 | -0.0210 | 0.0145 | 0.0239 |
| *RH_p10p* | -0.0127 | 0.0005 | 0.2700 | -0.0542 | -0.0472 | -0.0124 | 0.0234 | 0.0292 |
| *RH_IFSp* | -0.0108 | 0.0005 | 0.2885 | -0.0531 | -0.0455 | -0.0112 | 0.0255 | 0.0353 |
| *LH_a32pr* | -0.0106 | 0.0005 | 0.2965 | -0.0531 | -0.0465 | -0.0103 | 0.0248 | 0.0348 |
| *LH_p10p* | -0.0096 | 0.0005 | 0.3170 | -0.0511 | -0.0448 | -0.0105 | 0.0265 | 0.0333 |
| *LH_LIPv* | -0.0046 | 0.0005 | 0.4100 | -0.0466 | -0.0403 | -0.0049 | 0.0310 | 0.0407 |
| *RH_LIPv* | -0.0037 | 0.0005 | 0.4245 | -0.0459 | -0.0382 | -0.0047 | 0.0323 | 0.0401 |
| *RH_OFC* | 0.0071 | 0.0005 | 0.6285 | -0.0366 | -0.0283 | 0.0071 | 0.0431 | 0.0506 |
| *RH_V4t* | 0.0076 | 0.0005 | 0.6395 | -0.0345 | -0.0287 | 0.0075 | 0.0440 | 0.0527 |
| *RH_pOFC* | 0.0099 | 0.0005 | 0.6620 | -0.0349 | -0.0272 | 0.0092 | 0.0470 | 0.0560 |
| *RH_s32* | 0.0111 | 0.0005 | 0.6965 | -0.0313 | -0.0253 | 0.0114 | 0.0483 | 0.0562 |
| *LH_10v* | 0.0142 | 0.0005 | 0.7405 | -0.0263 | -0.0195 | 0.0136 | 0.0504 | 0.0585 |
| *RH_10v* | 0.0145 | 0.0005 | 0.7435 | -0.0297 | -0.0224 | 0.0144 | 0.0529 | 0.0606 |

| | | | | | | | | |
|---|---|---|---|---|---|---|---|---|
| *LH_V4* | 0.0158 | 0.0005 | 0.7710 | -0.0266 | -0.0196 | 0.0150 | 0.0530 | 0.0608 |
| *LH_s32* | 0.0170 | 0.0005 | 0.7730 | -0.0244 | -0.0176 | 0.0168 | 0.0544 | 0.0595 |
| *RH_V4* | 0.0171 | 0.0005 | 0.7920 | -0.0239 | -0.0180 | 0.0162 | 0.0538 | 0.0641 |
| *LH_OFC* | 0.0184 | 0.0005 | 0.8005 | -0.0250 | -0.0187 | 0.0179 | 0.0556 | 0.0632 |
| *LH_4* | 0.0190 | 0.0005 | 0.8130 | -0.0236 | -0.0162 | 0.0195 | 0.0548 | 0.0627 |
| *RH_4* | 0.0202 | 0.0005 | 0.8295 | -0.0210 | -0.0137 | 0.0201 | 0.0555 | 0.0633 |
| *LH_pOFC* | 0.0218 | 0.0005 | 0.8335 | -0.0246 | -0.0163 | 0.0222 | 0.0596 | 0.0678 |
| *LH_V4t* | 0.0236 | 0.0005 | 0.8580 | -0.0191 | -0.0124 | 0.0231 | 0.0605 | 0.0695 |
| ***Subcortical Regions*** | | | | | | | | |
| *RH_Caudate* | -0.0166 | 0.0005 | 0.2270 | -0.0588 | -0.0528 | -0.0174 | 0.0199 | 0.0302 |
| *RH_Pallidum* | -0.0096 | 0.0005 | 0.3240 | -0.0509 | -0.0444 | -0.0101 | 0.0254 | 0.0344 |
| *LH_Caudate* | -0.0048 | 0.0005 | 0.4000 | -0.0456 | -0.0395 | -0.0053 | 0.0302 | 0.0382 |
| *LH_Pallidum* | -0.0022 | 0.0005 | 0.4555 | -0.0452 | -0.0395 | -0.0023 | 0.0351 | 0.0426 |
| *LH_Putamen* | 0.0044 | 0.0005 | 0.5715 | -0.0381 | -0.0313 | 0.0040 | 0.0402 | 0.0471 |
| *RH_Putamen* | 0.0067 | 0.0005 | 0.6335 | -0.0362 | -0.0301 | 0.0063 | 0.0426 | 0.0509 |
| *LH_Accumbens* | 0.0192 | 0.0005 | 0.8030 | -0.0244 | -0.0172 | 0.0194 | 0.0569 | 0.0648 |
| *RH_Amygdala* | 0.0214 | 0.0005 | 0.8270 | -0.0211 | -0.0148 | 0.0204 | 0.0593 | 0.0671 |
| *LH_Amygdala* | 0.0219 | 0.0005 | 0.8355 | -0.0238 | -0.0161 | 0.0214 | 0.0628 | 0.0719 |
| *RH_Accumbens* | 0.0318 | 0.0005 | 0.9225 | -0.0121 | -0.0046 | 0.0313 | 0.0704 | 0.0793 |

**Supplementary Table 6.** Summary table for Future Expected Value (FEV) Bayesian Multilevel Model. Blue indicates regions with *P+* ≤ 0.15 and red indicates *P+* ≥ 0.85.

| ROI | *Mean* | *SE* | *P+* | *0.25* | *0.05* | *0.50* | *0.95* | *0.975* |
|---|---|---|---|---|---|---|---|---|
| ***Cortical Regions*** | | | | | | | | |
| *RH_pOFC* | -0.0144 | 0.0010 | 0.5020 | -0.1231 | -0.1069 | 0.0002 | 0.0397 | 0.0475 |
| *LH_pOFC* | 0.0016 | 0.0007 | 0.5975 | -0.0694 | -0.0566 | 0.0073 | 0.0417 | 0.0471 |
| *LH_OFC* | 0.0054 | 0.0006 | 0.6395 | -0.0530 | -0.0419 | 0.0090 | 0.0426 | 0.0482 |
| *RH_10v* | 0.0200 | 0.0004 | 0.8645 | -0.0202 | -0.0125 | 0.0213 | 0.0478 | 0.0530 |
| *RH_OFC* | 0.0199 | 0.0004 | 0.8680 | -0.0213 | -0.0127 | 0.0206 | 0.0495 | 0.0549 |
| *LH_4* | 0.0205 | 0.0004 | 0.8835 | -0.0162 | -0.0088 | 0.0212 | 0.0470 | 0.0539 |
| *LH_10v* | 0.0222 | 0.0004 | 0.8915 | -0.0135 | -0.0088 | 0.0220 | 0.0514 | 0.0575 |
| *RH_4* | 0.0216 | 0.0004 | 0.8925 | -0.0145 | -0.0089 | 0.0224 | 0.0494 | 0.0536 |
| *LH_IFSp* | 0.0247 | 0.0004 | 0.9235 | -0.0098 | -0.0038 | 0.0252 | 0.0517 | 0.0573 |
| *RH_V4* | 0.0277 | 0.0004 | 0.9455 | -0.0080 | -0.0005 | 0.0282 | 0.0546 | 0.0596 |
| *RH_s32* | 0.0292 | 0.0004 | 0.9520 | -0.0050 | 0.0007 | 0.0289 | 0.0568 | 0.0634 |
| *LH_s32* | 0.0287 | 0.0004 | 0.9525 | -0.0054 | 0.0003 | 0.0286 | 0.0557 | 0.0617 |
| *LH_LIPv* | 0.0302 | 0.0004 | 0.9555 | -0.0043 | 0.0018 | 0.0305 | 0.0576 | 0.0642 |
| *RH_p10p* | 0.0286 | 0.0004 | 0.9560 | -0.0052 | 0.0012 | 0.0286 | 0.0564 | 0.0619 |
| *LH_p9-46v* | 0.0298 | 0.0004 | 0.9645 | -0.0043 | 0.0033 | 0.0301 | 0.0577 | 0.0629 |
| *LH_V4* | 0.0313 | 0.0004 | 0.9645 | -0.0037 | 0.0033 | 0.0312 | 0.0592 | 0.0661 |

| | | | | | | | | |
|---|---|---|---|---|---|---|---|---|
| *RH_LIPv* | 0.0307 | 0.0004 | 0.9645 | -0.0032 | 0.0043 | 0.0313 | 0.0558 | 0.0621 |
| *LH_p10p* | 0.0311 | 0.0004 | 0.9680 | -0.0018 | 0.0040 | 0.0308 | 0.0586 | 0.0643 |
| *RH_p9-46v* | 0.0313 | 0.0004 | 0.9680 | -0.0022 | 0.0037 | 0.0310 | 0.0580 | 0.0640 |
| *RH_V4t* | 0.0310 | 0.0004 | 0.9680 | -0.0017 | 0.0036 | 0.0311 | 0.0586 | 0.0649 |
| *LH_V4t* | 0.0320 | 0.0004 | 0.9695 | -0.0017 | 0.0046 | 0.0317 | 0.0592 | 0.0643 |
| *RH_IFSp* | 0.0317 | 0.0004 | 0.9705 | -0.0013 | 0.0055 | 0.0322 | 0.0577 | 0.0632 |
| *RH_a32pr* | 0.0335 | 0.0004 | 0.9735 | -0.0002 | 0.0057 | 0.0332 | 0.0612 | 0.0673 |
| *LH_a32pr* | 0.0328 | 0.0004 | 0.9760 | 0.0004 | 0.0066 | 0.0328 | 0.0595 | 0.0651 |
| ***Subcortical Regions*** | | | | | | | | |
| *LH_Accumbens* | -0.0253 | 0.0009 | 0.3235 | -0.1012 | -0.0926 | -0.0197 | 0.0305 | 0.0370 |
| *RH_Accumbens* | 0.0021 | 0.0006 | 0.5710 | -0.0544 | -0.0449 | 0.0043 | 0.0401 | 0.0465 |
| *LH_Amygdala* | 0.0209 | 0.0004 | 0.8780 | -0.0173 | -0.0103 | 0.0213 | 0.0505 | 0.0559 |
| *LH_Putamen* | 0.0229 | 0.0004 | 0.9010 | -0.0113 | -0.0066 | 0.0234 | 0.0494 | 0.0557 |
| *RH_Putamen* | 0.0232 | 0.0004 | 0.9145 | -0.0143 | -0.0072 | 0.0235 | 0.0513 | 0.0561 |
| *RH_Amygdala* | 0.0259 | 0.0004 | 0.9285 | -0.0099 | -0.0029 | 0.0263 | 0.0528 | 0.0583 |
| *LH_Caudate* | 0.0280 | 0.0004 | 0.9525 | -0.0055 | 0.0002 | 0.0282 | 0.0557 | 0.0609 |
| *LH_Pallidum* | 0.0284 | 0.0004 | 0.9565 | -0.0051 | 0.0013 | 0.0287 | 0.0557 | 0.0611 |
| *RH_Caudate* | 0.0289 | 0.0004 | 0.9565 | -0.0047 | 0.0016 | 0.0291 | 0.0553 | 0.0610 |
| *RH_Pallidum* | 0.0335 | 0.0004 | 0.9770 | 0.0005 | 0.0071 | 0.0340 | 0.0612 | 0.0671 |

**Supplementary Table 7.** Summary table for Reward Prediction Error (RPE) Bayesian Multilevel Model. Blue indicates regions with *P+* ≤ 0.15 and red indicates *P+* ≥ 0.85.

| ROI | *Mean* | *SE* | *P+* | *0.25* | *0.05* | *0.50* | *0.95* | *0.975* |
|---|---|---|---|---|---|---|---|---|
| ***Cortical Regions*** | | | | | | | | |
| *LH_V4* | -0.1444 | 0.0006 | 0.0000 | -0.1940 | -0.1859 | -0.1450 | -0.1011 | -0.0934 |
| *RH_V4* | -0.1401 | 0.0006 | 0.0000 | -0.1916 | -0.1833 | -0.1394 | -0.0984 | -0.0911 |
| *RH_V4t* | -0.0606 | 0.0005 | 0.0050 | -0.1082 | -0.0998 | -0.0607 | -0.0225 | -0.0145 |
| *LH_LIPv* | -0.0364 | 0.0005 | 0.0620 | -0.0827 | -0.0763 | -0.0358 | 0.0027 | 0.0098 |
| *RH_LIPv* | -0.0317 | 0.0005 | 0.0830 | -0.0776 | -0.0700 | -0.0315 | 0.0060 | 0.0128 |
| *RH_a32pr* | -0.0226 | 0.0005 | 0.1670 | -0.0678 | -0.0601 | -0.0222 | 0.0136 | 0.0209 |
| *LH_a32pr* | -0.0072 | 0.0005 | 0.3845 | -0.0508 | -0.0449 | -0.0074 | 0.0319 | 0.0403 |
| *LH_V4t* | -0.0050 | 0.0005 | 0.4115 | -0.0495 | -0.0425 | -0.0055 | 0.0327 | 0.0398 |
| *RH_p9-46v* | 0.0042 | 0.0005 | 0.5770 | -0.0400 | -0.0331 | 0.0041 | 0.0401 | 0.0478 |
| *RH_IFSp* | 0.0228 | 0.0005 | 0.8565 | -0.0200 | -0.0126 | 0.0226 | 0.0589 | 0.0670 |
| *LH_p9-46v* | 0.0261 | 0.0005 | 0.8755 | -0.0173 | -0.0095 | 0.0260 | 0.0631 | 0.0695 |
| *LH_IFSp* | 0.0400 | 0.0005 | 0.9525 | -0.0069 | 0.0011 | 0.0405 | 0.0777 | 0.0829 |
| *RH_pOFC* | 0.0470 | 0.0006 | 0.9615 | -0.0054 | 0.0032 | 0.0464 | 0.0932 | 0.1007 |
| *RH_OFC* | 0.0573 | 0.0005 | 0.9905 | 0.0086 | 0.0168 | 0.0568 | 0.0963 | 0.1036 |
| *LH_4* | 0.0567 | 0.0005 | 0.9935 | 0.0140 | 0.0217 | 0.0565 | 0.0934 | 0.1001 |
| *RH_p10p* | 0.0600 | 0.0005 | 0.9940 | 0.0142 | 0.0212 | 0.0601 | 0.0972 | 0.1042 |

| | | | | | | | | |
|---|---|---|---|---|---|---|---|---|
| *LH_p10p* | 0.0636 | 0.0005 | 0.9970 | 0.0170 | 0.0251 | 0.0635 | 0.1029 | 0.1098 |
| *LH_10v* | 0.1526 | 0.0006 | 1.0000 | 0.1016 | 0.1102 | 0.1535 | 0.1936 | 0.2015 |
| *LH_OFC* | 0.1081 | 0.0006 | 1.0000 | 0.0611 | 0.0677 | 0.1076 | 0.1490 | 0.1595 |
| *LH_pOFC* | 0.0967 | 0.0006 | 1.0000 | 0.0444 | 0.0517 | 0.0971 | 0.1402 | 0.1493 |
| *LH_s32* | 0.1507 | 0.0005 | 1.0000 | 0.1036 | 0.1120 | 0.1507 | 0.1893 | 0.1956 |
| *RH_10v* | 0.1614 | 0.0006 | 1.0000 | 0.1148 | 0.1219 | 0.1610 | 0.2024 | 0.2093 |
| *RH_4* | 0.0932 | 0.0005 | 1.0000 | 0.0488 | 0.0555 | 0.0932 | 0.1292 | 0.1377 |
| *RH_s32* | 0.1462 | 0.0005 | 1.0000 | 0.0993 | 0.1080 | 0.1457 | 0.1861 | 0.1943 |
| ***Subcortical Regions*** | | | | | | | | |
| *LH_Pallidum* | 0.0127 | 0.0005 | 0.7300 | -0.0299 | -0.0234 | 0.0134 | 0.0484 | 0.0556 |
| *RH_Pallidum* | 0.0415 | 0.0005 | 0.9775 | 0.0019 | 0.0084 | 0.0409 | 0.0769 | 0.0822 |
| *LH_Accumbens* | 0.2700 | 0.0006 | 1.0000 | 0.2178 | 0.2274 | 0.2705 | 0.3128 | 0.3209 |
| *LH_Amygdala* | 0.0858 | 0.0006 | 1.0000 | 0.0362 | 0.0436 | 0.0856 | 0.1271 | 0.1354 |
| *LH_Caudate* | 0.0896 | 0.0005 | 1.0000 | 0.0452 | 0.0521 | 0.0891 | 0.1258 | 0.1330 |
| *LH_Putamen* | 0.0997 | 0.0005 | 1.0000 | 0.0583 | 0.0647 | 0.0993 | 0.1379 | 0.1448 |
| *RH_Accumbens* | 0.2775 | 0.0006 | 1.0000 | 0.2273 | 0.2350 | 0.2775 | 0.3186 | 0.3273 |
| *RH_Amygdala* | 0.1207 | 0.0005 | 1.0000 | 0.0750 | 0.0813 | 0.1212 | 0.1595 | 0.1680 |
| *RH_Caudate* | 0.0910 | 0.0005 | 1.0000 | 0.0462 | 0.0528 | 0.0905 | 0.1289 | 0.1368 |
| *RH_Putamen* | 0.1078 | 0.0005 | 1.0000 | 0.0659 | 0.0722 | 0.1080 | 0.1441 | 0.1505 |
